# Assembly and performance of a cholera RDT prototype that detects both *Vibrio cholerae* and associated bacteriophage as a proxy for pathogen detection

**DOI:** 10.1101/2024.08.23.609438

**Authors:** Md. Abu Sayeed, Imrul Kayes Nabil, Piyash Bhattacharjee, Md. Shawkat Hossain, Noor Jahan Akter, Romana Akter, Karen L Kelley, Mahbubul Karim, Yasmin Ara Begum, Taufiqur Rahman Bhuiyan, Firdausi Qadri, Ashraful Islam Khan, Eric J Nelson

## Abstract

**Introduction:** Cholera rapid diagnostic tests (RDTs) are vulnerable to virulent bacteriophage predation. We hypothesized that an enhanced cholera RDT that detects the common virulent bacteriophage ICP1 might serve as a proxy for pathogen detection. We previously developed a monoclonal antibody (mAb) to the ICP1 major capsid protein. Our objective herein was to design and assemble a first-of-its-kind RDT that detects both a bacterial pathogen (*Vibrio cholerae*) and associated virulent bacteriophage (ICP1).

**Method:** Candidate mAbs were expanded to increase design options and evaluated by immunological assays (ELISA; western blot). A subset of mAbs were selected for gold conjugation and printing on the RDT. The limit of detection (LOD) of prototype RDTs were determined in diarrheal stools with the addition of ICP1.

**Results:** Three mAb candidates were developed and evaluated for the capsid decoration protein (ORF123) and tail fiber protein (ORF93), and the prior mAb for the major capsid protein (ORF122). A single mAb sandwich RDT prototype for ORF122 was able to detect ICP1; RDTs with mAbs to ORF123 and ORF93 failed to detect ICP1 in single or dual sandwich configurations. Biologically meaningful LODs for ICP1 were achieved only after boiling the stool with ICP1; analysis by electron microscopy suggested increased epitope availability after boiling.

**Conclusion:** In this study, we demonstrate a proof of concept for a functional RDT that can detect both the primary pathogen and a common virulent bacteriophage as a proxy for pathogen detection. Further optimization is required before scaled production and implementation.

## INTRODUCTION

Cholera is an ancient diarrheal disease yet today remains a global public health problem, especially in Asia and Africa (1). Globally, cholera cases are under reported; there are at least 1.3 to 4.0 million cases and more than 20,000 deaths each year (2). In 2017, The World Health Organization (WHO) Global Task Force on Cholera Control (GTFCC) launched the "Ending Cholera: A Global Roadmap to 2030". One objective of the roadmap was to reduce cholera mortality by 90% by 2030 (3). Despite this ambitious goal, over 30 countries battled outbreaks in 2024 and the WHO declared a level-three emergency which is their highest level (4).

Cholera surveillance and early outbreak detection are essential components of the GTFCC road map (3). Cholera surveillance allows for the mapping of cholera hotspots. Early outbreak detection within these hotspots enables effective interventions, including reactive vaccination, rehydration points and aggressive hygiene/sanitation (3, 5-7). Microbial culture and quantitative polymerase chain reaction (PCR/qPCR) are conventional approaches for surveillance and diagnosis (8, 9). However, these gold-standards are time-consuming, require well-trained staff and demand ready access to laboratories. As a result, access to cholera diagnostics is often limited in resource-poor settings where cholera outbreaks occur. This limitation highlights the demand for a simple, accurate, and inexpensive RDT that requires little training (10, 11).

Approaches to lateral flow assay-based RDTs take several formats (12, 13). Commercialized cholera RDTs use a sandwich immunoassay format (14). The sandwich format can be designed with a single mAb or two different antibodies to perform the labeling and capture steps (10, 15). Most cholera RDTs use anti-*V. cholerae* LPS mAb to both label and capture the target in this configuration, a mobile mAb is used to gold-label the target in the sample pad and then fixed mAb captures the target *V. cholerae* at the test line (10, 14, 16-18).

Despite the development, evaluation and commercialization of several cholera RDTs, their scope of use is restricted to epidemiologic applications alone because of inconsistent performance, especially in field settings (9, 19). We previously showed that RDT sensitivity was compromised by the virulent ICP1 bacteriophage which is common in both Asia and Africa (8); ICP1 is specific for *V. cholerae* and uses LPS as its receptor (20, 21). To increase RDT sensitivity in the context of ICP1 phage predation, we developed an anti-ICP1 mAb against ICP1 major head protein to detect ICP1 as a proxy for *V. cholerae* detection (22). In this study, we sought to expand potential RDT design configurations by developing additional mAbs that target ICP1 putative tail fibers and a head decoration protein. Using these mAbs, we developed both single and double mAbs-based RDT prototypes. The performance of the novel RDT configurations were evaluated in diarrheal stool samples spiked with ICP1 as critical steps towards a large clinical diagnostic study.

## METHODS

### Ethics Statement

The clinical samples used to develop the RDT were obtained through prior studies approved by the Research Review Committee (RRC) and the Ethical Review Committee (ERC) of the International Centre for Diarrhoeal Disease Research, Bangladesh (icddr,b), and the Institutional Review Boards (IRBs) of the Institute of Epidemiology, Disease Control and Research (IEDCR), and the University of Florida; the recruitment, consent, enrollment, and procedures were described previously (23, 24).

### Bacterial Strains and Phage Stock Preparation

We used *V. cholerae* O1 strain HC1037 (provided by Dr. Andrew Camilli, Tufts University) to make high-titer vibriophages ICP1, ICP2, and ICP3 using the previously described method (22, 25). This strain was selected because it naturally lacks K139 prophage and is sensitive to all three lytic phages. The bacterial strain was grown to a mid-log phase in Luria-Bertani (LB) broth at 37°C in a shaking incubator. The bacterial culture was inoculated with the phage for 4-6 hours. The phage stock was then prepared by two times polyethylene glycol (PEG) precipitation and stored in phage80 buffer. The high titer phage preparation was enumerated as PFU/mL on 0.35% soft agar media using standard methods (22). We prepared formalin-killed *V. cholerae* whole cell (VCWC) by treating mid-log bacterial culture with 0.5% formalin overnight at room temperature (RT).

### ICP1 recombinant antigen preparation

Two putative tail fiber proteins (ORF93 and ORF69) and a head decoration protein (ORF123) were cloned, expressed, and purified following the same methods used in our previous study on the ICP1 major head protein, ORF122 purification (22). Briefly, we cloned the targets into the pET16b vector (Novagen) using two restriction enzymes, NdeI and XhoI. We then transformed *Escherichia coli* (*E. coli*) BL21 (Novagen, Sigma-Aldrich) with the recombinant pET16b vector and induced the expression of His-tagged fusion proteins with Isopropyl β-d-1-thiogalactopyranoside. The recombinant proteins were then purified using Bugbuster reagent and His·Bind purification kit (Novagen) following the manufacturer’s user protocol. The concentration of the purified proteins was determined by standard Bio-Rad protein assay (26).

### Monoclonal antibody production

Hybridoma and cell culture techniques were contracted to ProMab Biotechnologies Inc. (Richmond, CA) to generate mAbs against the recombinant proteins (22). We received culture supernatants from 10 hybridoma clones per target from the vendor. After screening the clones (below), scaled production of the selected clones used both cell culture methods and the mouse ascites model (10).

### Indirect ELISA

Hybridoma clone culture supernatants were screened by an indirect ELISA (22). We coated Nunc MaxiSorp plates with ICP1 (10^8^ PFU/well), ICP2 (10^7^ PFU/well), ICP3 (10^8^ PFU/well), VCWC (10^6^ CFU/well), recombinant proteins (200 ng/well), and Bovine serum albumin (BSA; T200 ng/well). The plates were blocked with 1% BSA-Phosphate buffered saline (PBS) and incubated with a given hybridoma clone supernatant at a 1:20 dilution at 37°C for 1 hour. After incubating with horseradish peroxidase-tagged goat anti-mouse IgG (Jackson ImmunoResearch; 1:1000 dilution), the plate was developed using a chromogenic substrate,1-Step Ultra TMB. The reaction was then stopped with 1N H_2_SO_4_ before measuring the absorbance at 450 nm using an ELISA plate reader (SYNERGYMx, BioTek). The absorbance represented the reactivity of culture supernatants to the coated antigens.

### Western blot assay

We prepared the antigens by boiling them with NuPAGE SDS sample buffer for 10 minutes. The antigens were electrophoresed on NuPAGE 4 to 12% Bis-Tris precast gel (ThermoFisher) and blotted on a 0.2 µm nitrocellulose membrane using the Trans-Blot turbo Transfer System (Bio-Rad) (22). After blocking with 5% skim milk in Tris-buffered saline (TBS), the membrane was incubated at RT with 1:200 diluted hybridoma clone culture supernatants for 1 hour. The membrane was then treated with a secondary antibody, alkaline phosphatase-conjugated goat anti-mouse IgG (1:5000 dilution) for 1 hour at RT. Finally, the membrane was developed using 5-bromo-4-chloro-3-indolyl-phosphate/nitro blue tetrazolium (BCIP/NBT) substrate, and the image of the membrane was taken by a gel imager (Geldoc; Bio-Rad).

### Colloidal Gold and Gold Conjugate Preparation

We boiled chloroauric acid (HAuCl4; 0.01%) with sodium citrate (0.024%) until the solution appeared a red wine color. Sodium citrate acted as a reducing agent, and this reduction process generated 20 nm colloidal gold (27). The solution was then filtered through a 0.2 µm filter before conjugation with the detection antibody. An aggregation test was used to optimize minimum protein concentration and optimum pH for gold conjugation. We conjugated gold particles with anti-ORF122, ORF123, and ORF93 mAbs at different pH and concentrations. We then added 10% NaCl to the conjugate solution for 10 minutes to perform the aggregation test. The absorbance at 520 nm, 580 nm, and 600 nm was measured to check the stability and polydispersity of the solution. We determined the optimum reaction conditions for gold conjugation (see below). After adding 20% BSA, the gold solution was centrifuged at 10000 rpm for 45 minutes at 4°C. The pellet was then resuspended in 1% BSA-0.002M Tris buffer and filtered in a 0.2 µm filter before use in the conjugate pad. The conjugation of *V. cholerae-*specific mAb (anti-VC LPS mAb) was described previously (10).

### RDT prototype assembly

We assembled two RDT protypes: ‘ICP1 RDT’ and ‘RDTplus’. To optimize ICP1 detection, we used the ICP1 RDT prototype in which we dispensed only one test line (anti-ICP1 ORF122/ORF123/ORF93 mAb; 1 mg/ml) on a nitrocellulose membrane (High flow plus 120 Membrane card; Millipore). For the ‘RDTplus’ prototype, we modified the existing Cholkit by dispensing two test lines with anti-ICP1 ORF122 mAb (1.0 mg/ml) and anti-VC LPS mAb (0.35 mg/ml), respectively, on the nitrocellulose membrane tagged at the middle of a backing card. For both prototypes, we dispensed the control line with goat anti-mouse IgG (1 mg/mL). The membrane was dried at 45°C for 90 minutes followed by blocking with 1% BSA-PBS for 20 minutes and again dried for 150 minutes. To prepare the conjugate pad, we soaked the glass fibers with anti-ICP1 ORF122/ORF123/ORF93 mAb-gold and anti-VC LPS mAb -gold conjugate (mobile detection antibodies) solution and air-dried them for two hours. For the ICP1 RDT prototype, we used only one type of conjugate pad (anti-ICP1 ORF122/123/93 mAb-gold), whereas two types of conjugate pads (anti-ICP1 ORF122 mAb-gold and anti-VC LPS mAb -gold) were used in RDT plus prototype. The conjugate pads were attached at the bottom edge of the nitrocellulose membrane. Another glass fiber sample pad was placed just below the conjugate pad in an overlapping manner. We then attached a cellulose fiber absorbent pad at the top edge of the nitrocellulose membrane to facilitate the sample flow through the RDT strip. We cut the backing card, assembled with all components, into 3 mm strips with a Guillotine cutter (CTS300 and ZQ2002).

### Sample Processing and Testing RDT Prototype

To optimize the performance of the RDT prototype, we processed samples with different physical and chemical treatments. We boiled high-titer ICP1 at different times to observe the effect of sample boiling time on RDT performance. In the stool spike assay, we evaluated RDT performance on ICP1 spiked stool samples that were prepared under different conditions and with different manipulations (pH, DMSO, dialysis, filtration, centrifugation). Samples were diluted in 0.2 M Tris-0.5 M NaCl-0.5% Tween in a microcentrifuge tube. We then dipped the RDT prototype strip into the samples for up to 30 minutes. The appearance of a red line for both the test line(s) and control line indicated a positive result (10).

### Electron microscopy

Phage were examined by transmission electron microscopy negative stain and immunogold electron microscopy. Glow-discharged 400 mesh carbon coated Formvar copper grids (Electron Microscopy Sciences, Hatfield, PA) were floated onto 5 µl of vibriophage suspension for 5 minutes. Excess solution was blotted with filter paper and placed onto a drop of 1% aqueous uranyl acetate for 30 seconds. The excess uranyl acetate was blotted dry and examined with a FEI Tecnai G2 Spirit Twin TEM (FEI Corp., Hillsboro, OR) and digital images were acquired with a Gatan UltraScan 2k x 2k camera and Digital Micrograph software (Gatan Inc., Pleasanton, CA). For immunogold labeling, Poly-L-Lysine (Sigma-Aldrich, St. Louis, MO) treated 400-mesh carbon coated Formvar nickel grids were floated onto 10 µl of vibriophage suspensions for 5 minutes. The samples were fixed and crosslinked to the poly-L-lysine grids with 2% paraformaldehyde in PBS and washed with PBS. The grids were floated on blocking agent (1% non-fat dry milk, 0.5% cold water fish skin gelatin, 0.01% Tween-20 in PBS) then incubated with mouse primary antibody. Negative controls were prepared by replacing primary antibody with PBS. Grids were washed in PBS and incubated with a 12 nm Colloidal Gold AffiniPure Goat Anti-Mouse IgG (1:20 dilution; Jackson ImmunoResearch Laboratories, West Grove, PA), washed in PBS, fixed with Trump’s fixative (Electron Microscopy Sciences, Hatfield, PA), and water washed. Once dried, the sample grid was floated on a 10 µl droplet of 1% aqueous uranyl acetate for 30 seconds, stain removed with filter paper, air dried and examined with a FEI Tecnai G2 Spirit Twin TEM (FEI Corp., Hillsboro, OR) and digital images were acquired with a Gatan UltraScan 2k x 2k camera and Digital Micrograph software (Gatan Inc., Pleasanton, CA).

### Statistical, Molecular and Bioinformatics Analysis

We used GraphPad Prism version 8 (GraphPad Software, Inc.) for data analysis, and graphical presentation. Bioinformatic analysis of target protein sequences from the published Bangladesh and the DRC ICP1 genome sequences (28, 29) were done by Geneious (Dotmatics). We also used polymerase chain reaction and Sanger sequencing to generate sequences from clinical samples collected from Bangladesh, the Democratic Republic of Congo (DRC), and Kenya using the primers listed below (Table S1). We then performed multiple sequence alignment (MSA) to explore the conservation of these sequences in phages isolated across different geographical regions. We used QIAGEN CLC software for the MSA.

## RESULTS

### Candidate ICP1 antigen selection, characterization, and development

To develop mAbs against ICP1, we targeted ICP1 structural proteins. We previously developed a mAb against the ICP1 major head protein (ORF122). We selected ORF122 protein because it was highly conserved among ICP1 stains collected at different time periods and geographical locations (22). To expand assembly options for the RDT prototype, we developed mAbs to additional structural targets. We characterized five additional putative ICP1 structural proteins using genomic sequences from Bangladesh (28) and the Democratic Republic of Congo (DRC) (29). These proteins showed 95-99% similarity at the amino acid level (Table S2). We next narrowed our candidate list to two putative tail proteins (ORF69 and ORF93) and the putative head decoration protein (ORF123) based on conservation. We performed MSA using the prior genomic sequences and sequences we generated by PCR/Sanger sequencing from clinical samples collected from Bangladesh, DRC and Kenya. The MSA demonstrated that across the geographical regions, ORF69, ORF93, and ORF123 sequences showed 98.5-100%, 94-100%, and 98.4-100% conservation at the amino acid sequence level, respectively (Figure S1; Table S3) and 99.5-100%, 90.4-100%, and 99.4-100% similarity at the nucleic acid level, respectively (Figure S2-S4; Table S3). A high level of conservation in the tail and head decoration proteins, irrespective of geographic site of isolation, supported their candidacy for mAb development. In addition to conservation, we sought mAbs that targeted distant sites in the phage anatomy to avoid epitope shielding (e.g., head vs tail). Hence, conserved proteins that targeted the tail fibers (ORF69 and ORF93) were selected for mAb development in addition to the head decoration protein ORF123 (see methods; Figure S5).

### Evaluation of ICP1 Reactive mAbs by immunoassays

To evaluate the new mAbs, we performed an indirect ELISA and screened the culture supernatants from the clones for their reactivity to PEG purified ICP1 (Figure 1). We found three ORF93-specific clone supernatants (ICP1ORF93_mAb CL1, CL6, and CL8) that were reactive against ICP1. Three ORF123-specific clone supernatants (ICP1ORF123_mAb CL14, CL15, and CL16) were also highly reactive to ICP1. However, none of the ORF69 clone supernatants were reactive to ICP1. All clone supernatants were non-reactive to our negative controls (ICP2, ICP3, VCWC, BSA). We next analyzed all ICP1 reactive clone supernatants by Western blot analysis (Figure S6). All ORF123-specific clone supernatants, ICP1ORF123_mAb CL14, CL15, and CL16, but not ORF95-specific clones, were able to detect PEG purified ICP1 on the Western blot membrane without cross-reactivity to the negative controls. Similar to the ELISA results, none of the ORF69-specific clone supernatants reacted to ICP1. ICP1ORF123_mAbCL14 and ICP1ORF93_mAb CL6 were selected for further analysis in RDT prototype development, hence forth referred to as ORF123 mAb and ORF93 mAb, respectively.

**Fig 1.**
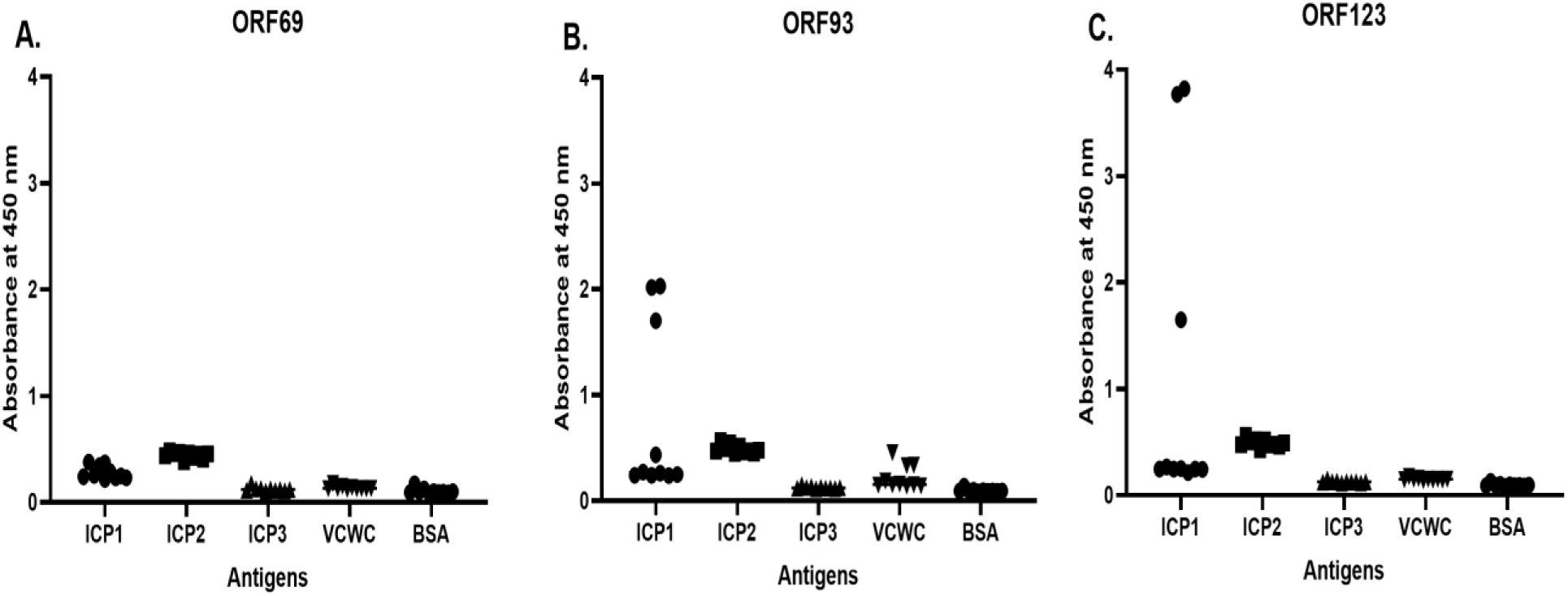
IgG antibody responses by ELISA in hybridoma clone culture supernatants derived from mice immunized with recombinant proteins: ICP1 tail fiber ORF69 (**A**), tail fiber ORF93 (**B**), and head decoration protein ORF123 (**C**). X axis represents the antigens assayed; VCWC = formalin-killed *V. cholerae* whole-cell, and bovine serum albumin = BSA. The Y axis is the absorbance at 450 nm read in SYNERGY Mx (BioTek) plate reader.

### Testing and optimization of the RDT prototype

#### Selection of labeling and capture antibodies for RDT prototyping

Based on the immune-assay results, we selected ORF123 and ORF93 mAbs in addition to the prior ORF122 mAb for developing a RDT prototype (22). We created a matrix strategy to generate an array of prototypes in which mAbs would serve as both labeling and capture antibodies. In addition, we allowed for both single and dual mAb sandwich formats. To prepare the labeling mAb for RDT prototype assembly, we conjugated ORF122, ORF123, and ORF93 specific mAbs with colloidal gold. Prior to conjugation, we determined the optimum conditions for mAb-gold conjugation. We found the optimum pH-9 and minimum mAb concentration of 20 µg/mL for conjugation for all mAbs except ORF93 (30 µg/mL), as the mAb-gold conjugates showed maximum stability and minimal polydispersity in these conditions (Figure S7). The high stability of the colloidal gold solution was represented by a high ratio of absorbance at 520 nm to 580 nm, whereas the minimal polydispersity was determined by the lowest absorbance ratio of the colloidal gold solution at 600 nm and 520 nm (27).

#### RDT detection of purified ICP1

None of the sandwich combinations were able to detect PEG purified ICP1. However, after boiling the ICP1 substrate, the ORF122mAb:ORF122mAb sandwich alone detected ICP1 (positive test line; positive control line). Therefore, we selected this RDT prototype for further analysis and development. We found an increase in test line intensity with increased duration of boiling the ICP1 substrate; maximal test line intensity was reached at 10 minutes of boiling (Figure 2A). Similar results were observed when we merged ORF122 mAb:ORF122 mAb sandwich RDT format with Cholkit to develop the ‘RDTplus’ prototype. The RDTplus prototype detected ICP1 after boiling the ICP1 test substrate for 10 min (Figure S8). The limit of detection of ICP1 was determined as 1.35×10^7^ PFU/mL when ICP1 prep was diluted in PBS (Figure 2B). This RDT prototype showed no cross-reactivity when we tested against *V. cholerae*, ICP2, and ICP3.

**Fig 2.**
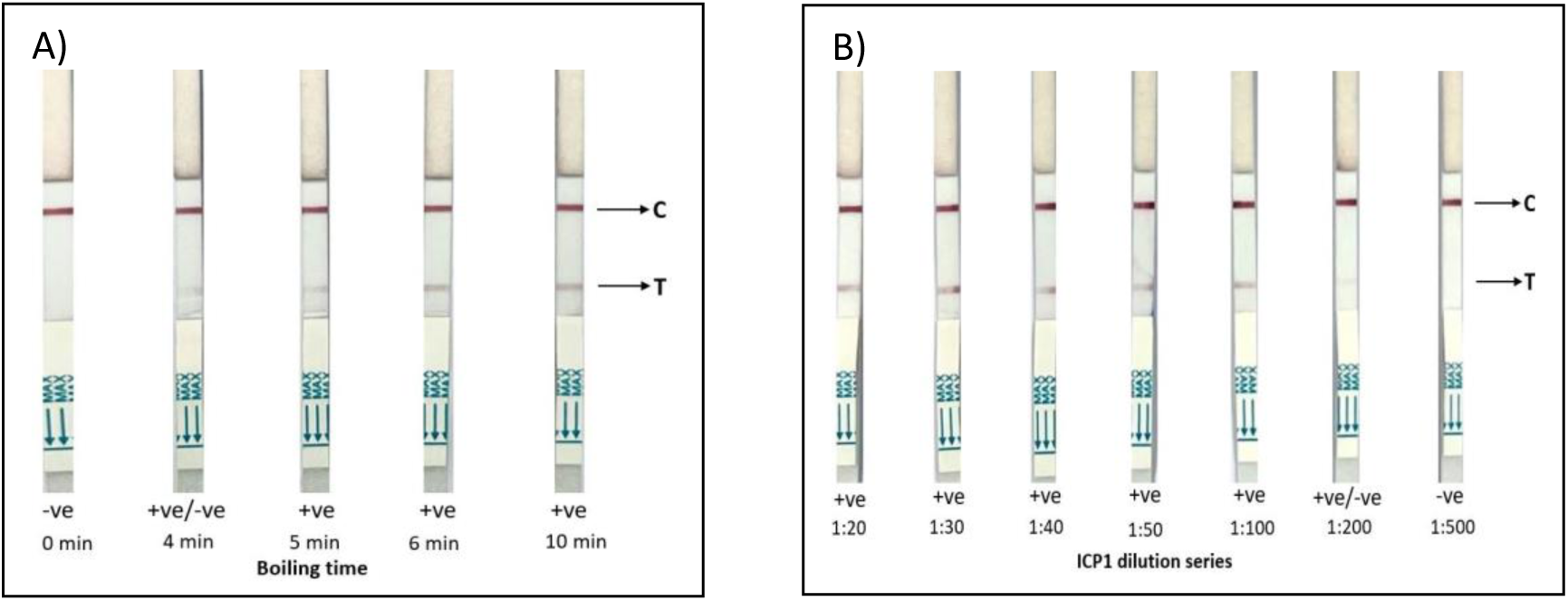
Testing RDT prototype for ICP1 detection using different boiling periods at a 1:10 dilution of high titer ICP1 (1×10^9^ PFU/mL) in 0.5X PBS (A), and then with dilutions ranging from 1:20 (5×10^7^ PFU/ml) to 1:500 (2×10^6^ PFU/ml) after 10 min boil (B). Here, C = control line, T= test line, -ve= negative result and +ve=positive result. Red lines indicate positive control or test line. Positive C line ensures the validity of the RDT prototype result.

#### RDT detection of ICP1 spiked in diarrheal stool matrix

To evaluate the ORF122 mAb:ORF122 mAb RDT prototype for diagnostic application, we generated mock stool samples with ICP1. We used a higher titer PEG purified ICP1 stock at 1.35×10^9^ PFU/mL to spike three *V. cholerae* and ICP1 negative stools (EN70, EN105, and EN122) collected in a prior study at a final biologically relevant concentration of 1.35×10^8^ PFU/mL (24). All three spiked stools samples tested positive for the ICP1 test line after 10 min of boiling (Figure 3); the intensity of the test line in spiked samples varied between samples. The intensity was lower compared to ICP1 diluted in PBS. The test line intensity moderately increased after the raw stool spiked with ICP1 was boiled and the time was increased to 20 or 30 minutes (Figure 3). The limit of detection was 3.4×10^7^ PFU/mL to 6.8×10^7^ PFU/mL after 20 min of boiling the substrate.

**Fig 3.**
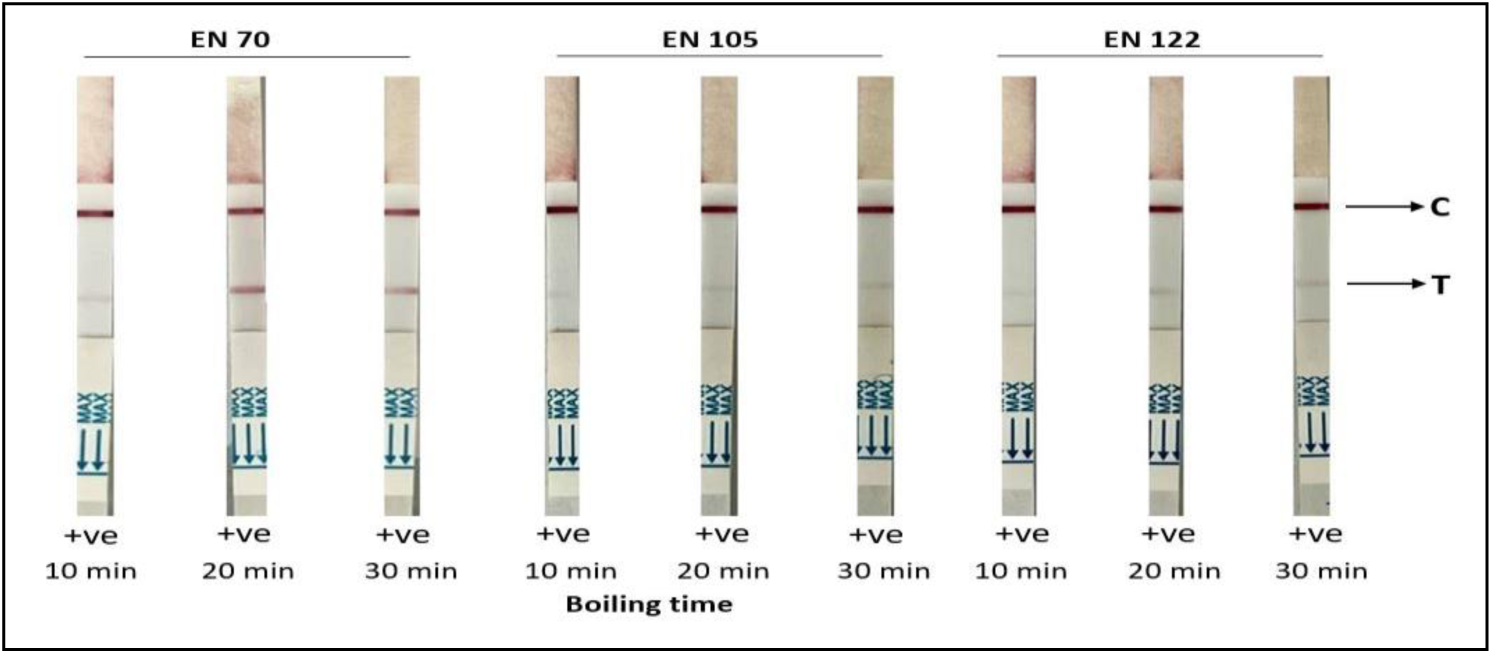
Evaluation of the RDT prototype for ICP1 detection in spiked diarrheal stools (EN 70, EN 105 and EN 122. High titer ICP1 (1×10^9^ PFU/mL) was spiked at 1:10 dilution and boiled at 95°C for 10, 20 and 30 minutes. Here, C = control line, T= test line, -ve= negative result (none) and +ve=positive result (all). Red lines indicate positive control or test line. Positive C line ensures the validity of the RDT prototype result.

We hypothesized that there was an inhibitory factor in the stool matrix that decreased the detection of ICP1. We tested this hypothesis using a previously obtained stool sample (30) that exhibited the strongest negative effect on RDT ICP1 detection (EN105). We first used dialysis to determine if the inhibitory factor (s) was a large vs a small molecule using a 10kD dialysis membrane; the inhibitory effect was identified in the large molecular fraction (not the dialysate). We explored if we could mitigate the inhibitory effect using techniques that would add minimal cost and effort to procedural steps for RDT workflows. We found DMSO treatment of the stool matrix showed a negative impact on RDT performance, potentially by interfering with the gold-conjugated detection antibody. Additional maneuvers that included altering the pH, filtration (0.2 μM), and centrifugation of the raw stool matrix did not increase the test line’s intensity.

### Determine the relative abundance of epitopes by EM analysis

We performed EM analysis to characterize how the anti-capsid (ORF122) mAb bound ICP1 (Figure 4). Without boiling and without immunogold labeling, ICP1 were associated with outer membrane vesicles (OMVs) which is consistent with prior studies (31). With immunogold labelling and no boiling, few gold-labeled anti-capsid mAb associated with ICP1. However, after 10 min and 20 min of the boiling the ICP1 substrate, the gold-labeled anti-capsid mAb abundantly associated with ICP1 remnants. Specifically, 12 nm gold particles decorated intact capsids and fragmented capsids with and without associated neck and tail fibers. These effects were most pronounced with 20 minutes of boiling the ICP1 substrate. In these preparations, non-specific/ background staining with the 12 nm gold particles was minimal.

**Fig 4.**
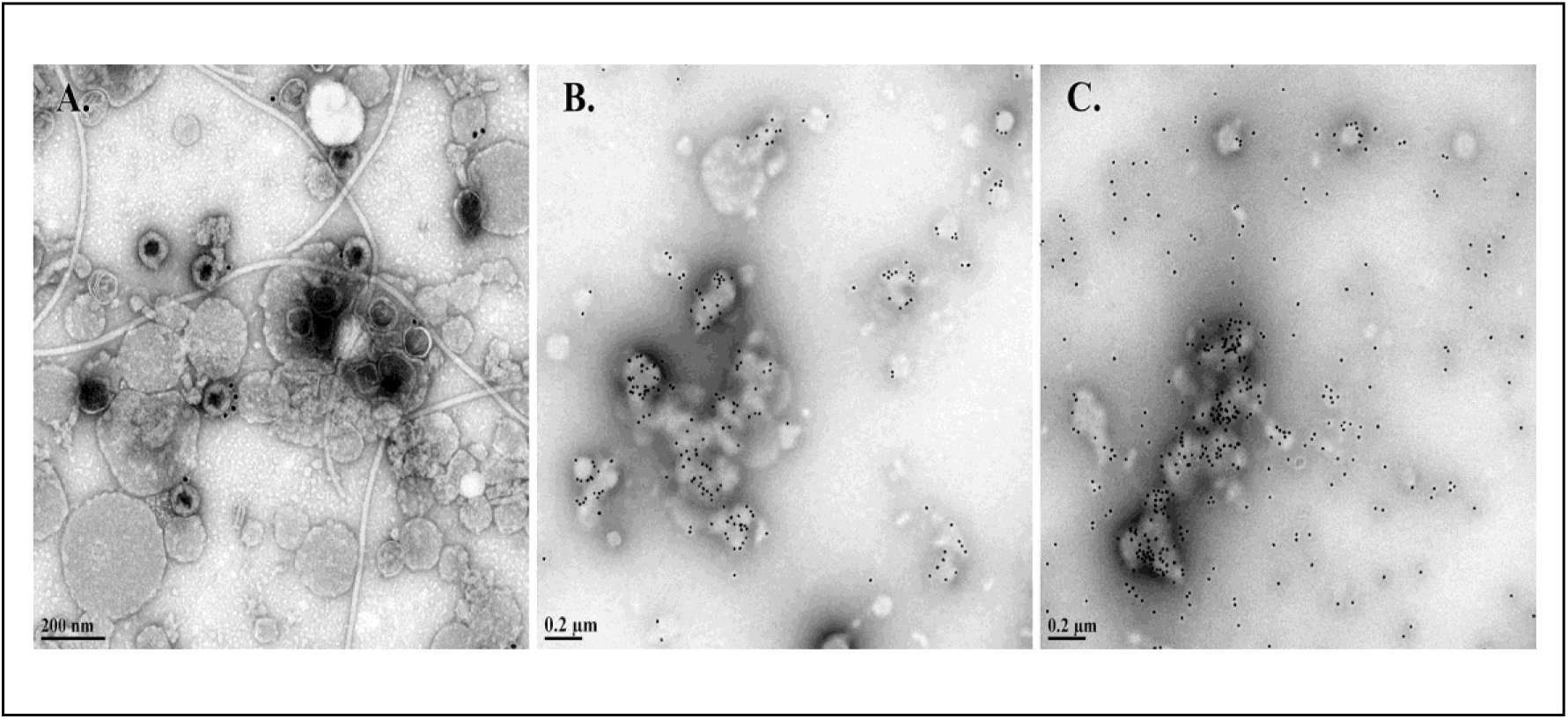
Electron micrographs of the ICP1 preparation (1×10^9^ PFU/mL) at a 1:10 dilution without boiling (**A**), with 10 min boiling **(B)** and with 20 min boiling **(C)** after immunolabeling with ICP1ORF122 mAb. Scale bars are imbedded in images.

## DISCUSSION

This study aimed to design and assemble a first-of-its-kind RDT that detects both a bacterial pathogen (*Vibrio cholerae*) and associated virulent bacteriophage (ICP1). After multiple iterations, the design with the most robust configuration was an RDT that included the prior single anti-LPS mAb to gold-label and capture the *V. cholerae* target, and now, a single mAb to the major capsid protein (ORF122) to gold-label and capture ICP1 phage particles. Biologically meaningful LODs for ICP1 were achieved however only after boiling stool with ICP1. The finding was supported by EM that suggested increased epitope availability after boiling. Therefore, we provide proof of concept for a functional RDT prototype (‘RDTplus’) that can detect a virulent bacteriophage as a proxy for pathogen detection, however further optimization is required before scaled production and implementation.

During the iterative design process, we first focused on published *in silico* analyses of the ICP1 putative structural proteins, including head and tail proteins (21). We previously demonstrated that the major head protein (ORF122) of ICP1 is immunogenic and can be used to generate a mAb against ICP1 (22). We selected two putative tail fiber proteins (ORF69 and ORF93) and one putative head decoration protein (ORF123) as immunogens for developing a second set of antibodies against ICP1. The rationale for these candidates was that they were highly conserved among ICP1 sequences in Asia and Africa which hopefully would convey durability in the context of high rates of evolutionary changes in both *V. cholerae* and associated virulent phages (21, 29, 32). In addition, we wanted structural protein candidates that were anatomically distant to avoid epitope shielding.

While performing immunoassays to screen the hybridoma clone culture supernatants, we found that three tail fiber (ORF93) clones and three head decoration protein (ORF123) clones were highly reactive and specific to ICP1. These findings are consistent with other phage immunogenicity studies in animal models. While the literature on phage immunobiology is scant, *Staphylococcal* bacteriophages induce specific antibody responses in mice against head and tail proteins (33), and the major head protein and head decoration protein of *E. coli* T4 phage found are highly immunogenic (34, 35). We analyzed these six ICP1 reactive clones by Western blot. All three tail fiber (ORF123) specific clones were able to detect ICP1, whereas the tail fiber (ORF93) specific clones could not detect ICP1; analyses were not performed to investigate the mechanistic failure of the ORF93 mAbs.

It was unexpected that the RDT prototypes in single or double mAb sandwich configurations were unable to detect ICP1. Potential reasons for this failure may be epitope saturation or epitope shielding which leaves limited unbound epitopes for capture at the test line. These challenges are common (36, 37); for example, a study on the therapeutic mAb daratumumab found that the mAb saturates the myeloma cell marker CD38 and interferes with the diagnostic CD38 antibodies (38). Despite this challenge, we found that boiling ICP1 substrate enabled a single mAb sandwich configuration (ORF122 mAb::ORF122 mAb) to detect ICP1 spiked into cholera stool at meaningful biologic concentrations. The EM analysis found increased gold-label binding to capsids and capsid fragments among samples boiled compared to samples not boiled. The EM approach could not resolve epitope location; we hypothesize the accessible epitopes may reside inside of the head structure and/or are shielded by proteins that must be degraded by boiling to expose the ORF122 epitope. Future research may be required to further optimize the RDT prototype. Alternatively, a new suite of mAbs to ORF122 protein could be generated and epitope location could be validated with cryo-electron microscopy (39, 40).

These findings need to be interpreted in the context of the study limitations. Firstly, the RDT prototype in its current configuration requires a boil step which impedes scalability. Secondly, the study presents detection of ICP1 spiked into cholera samples and not detection in stool samples with native ICP1 which requires a prospective clinical study. Lastly, the spatial positions of the accessible epitopes need to be analyzed by cryo-electron microscopy to further understand the limitations with the current design. Despite these limitations, the detection of ICP1 spiked into cholera stool represents a critical proof of concept for RDT development to detect bacterial pathogens directly or indirectly by detecting pathogen-specific bacteriophage as a proxy.

## CONCLUSIONS

Bacterial diagnostics are vulnerable to virulent bacteriophage predation which can degrade pathogen nucleic acid within minutes after injection and lyse a high percentage of the pathogen population within a few generations. We have documented aspects of this problem in cholera and seek solutions for both clinical, environmental and laboratory settings. In this study, we demonstrate a proof of concept for a fieldable RDT that can detect both the primary pathogen as well as a common virulent bacteriophage as a proxy for pathogen detection. Further optimization is required before scaled production and implementation.

## Acknowledgements

We thank the patients for participating in the studies from which the clinical samples were obtained in the prior studies, as well as the research teams at Incepta and the icddr,b who helped to make this work possible. We are grateful to Brittney Johnson, Randy Autrey, Krista Berquist for their administrative expertise, as well as the UF Emerging Pathogens Institute and UF the Department of Pediatrics for providing vital infrastructure. We are grateful to Abdul Muktadir (Incepta Chairperson) and Hasneen Muktadir (Incepta Chairperson) for their support of this initiative and commitment to initiatives at Incepta that aim to improve the health for all persons in Bangladesh and globally. We thank Mary Gragg and Quinnton Cooper at the UF ICBR Electron Microscopy facility (RRID:SCR_019146) for their assistance during the electron microscopy experiment.

## Data availability

Data analyzed are presented within the manuscript and online supplementary material.

## Disclaimer

The funders had no role in the study design, data collection and analysis, decision to publish, or preparation of the manuscript.

## SUPPLEMENTARY MATERIALS

**Fig S1.**
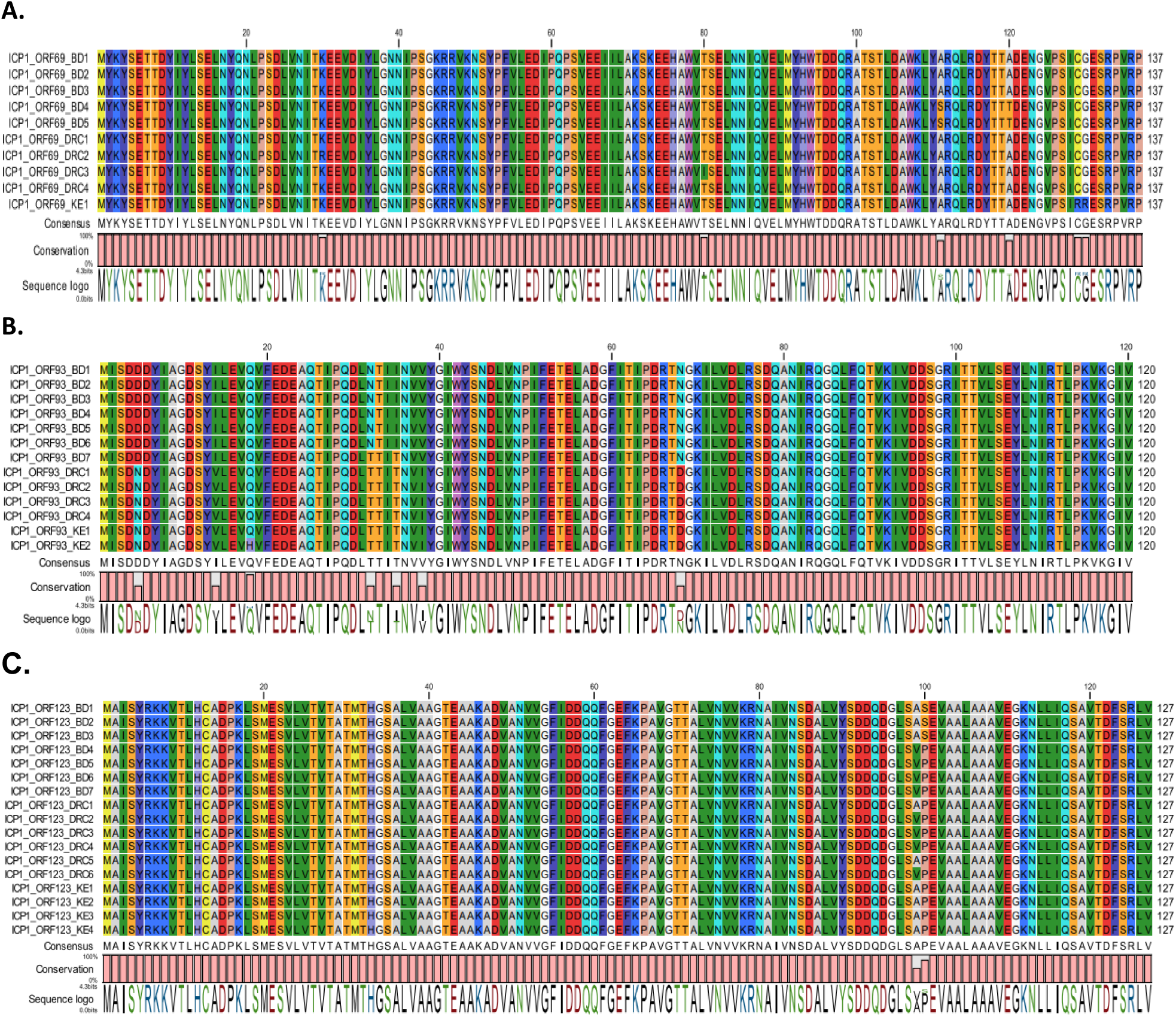
Amino acid alignment of ICP1 tail fiber ORF69 (**A**), tail fiber ORF93 (**B**), and capsid decoration protein ORF123 (**C**) from Bangladesh (BD), Democratic Republic of Congo (DRC), and Kenya (KE). Sequences were obtained by PCR amplification and sequencing of clinical samples. Data visualizations prepared in QIAGEN CLC.

**Fig S2.**
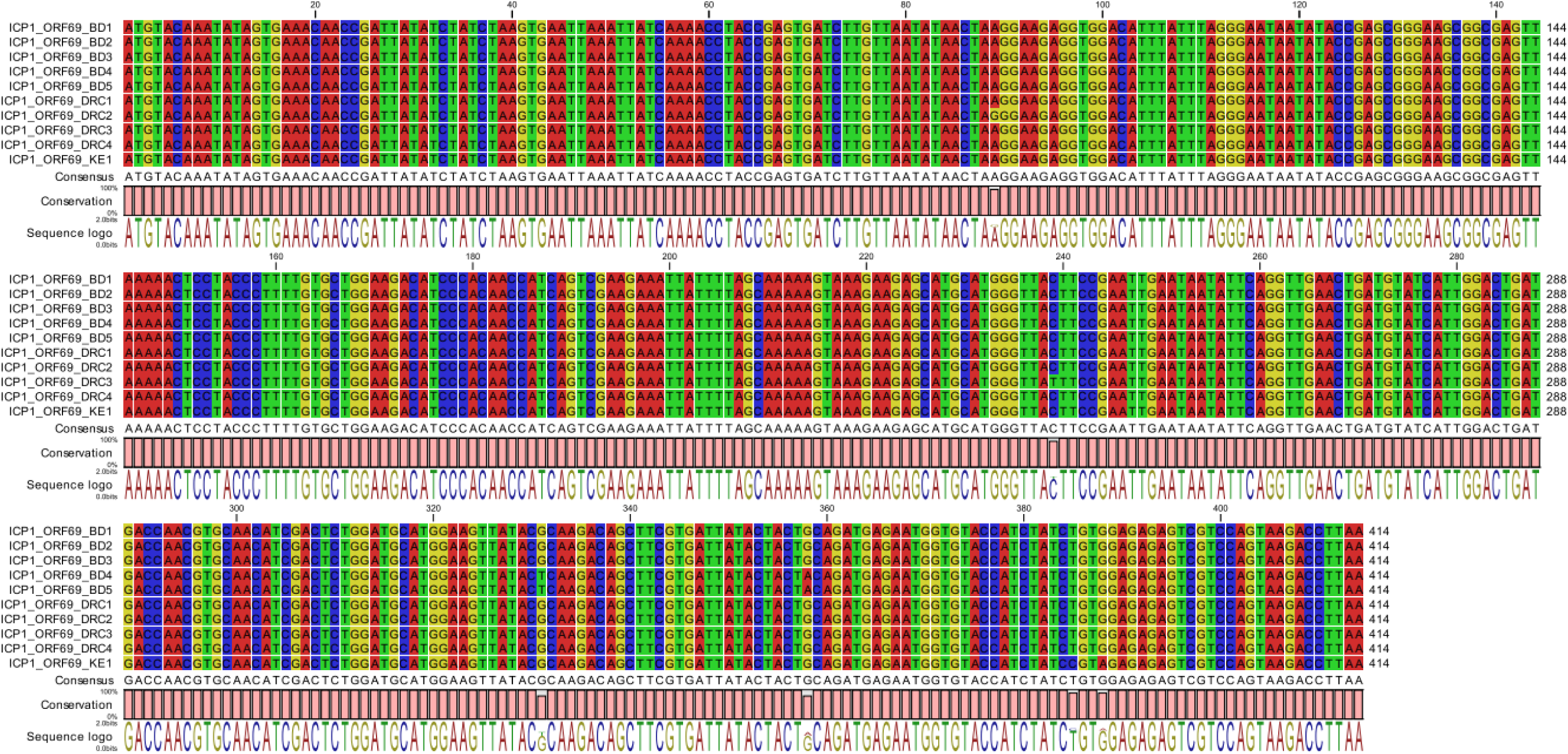
Nucleic acid alignment of ICP1 tail fiber ORF69 from Bangladesh (BD), Democratic Republic of Congo (DRC), and Kenya (KE). Sequences were obtained by PCR amplification and sequencing of clinical samples. Data visualizations prepared in QIAGEN CLC software.

**Fig S3.**
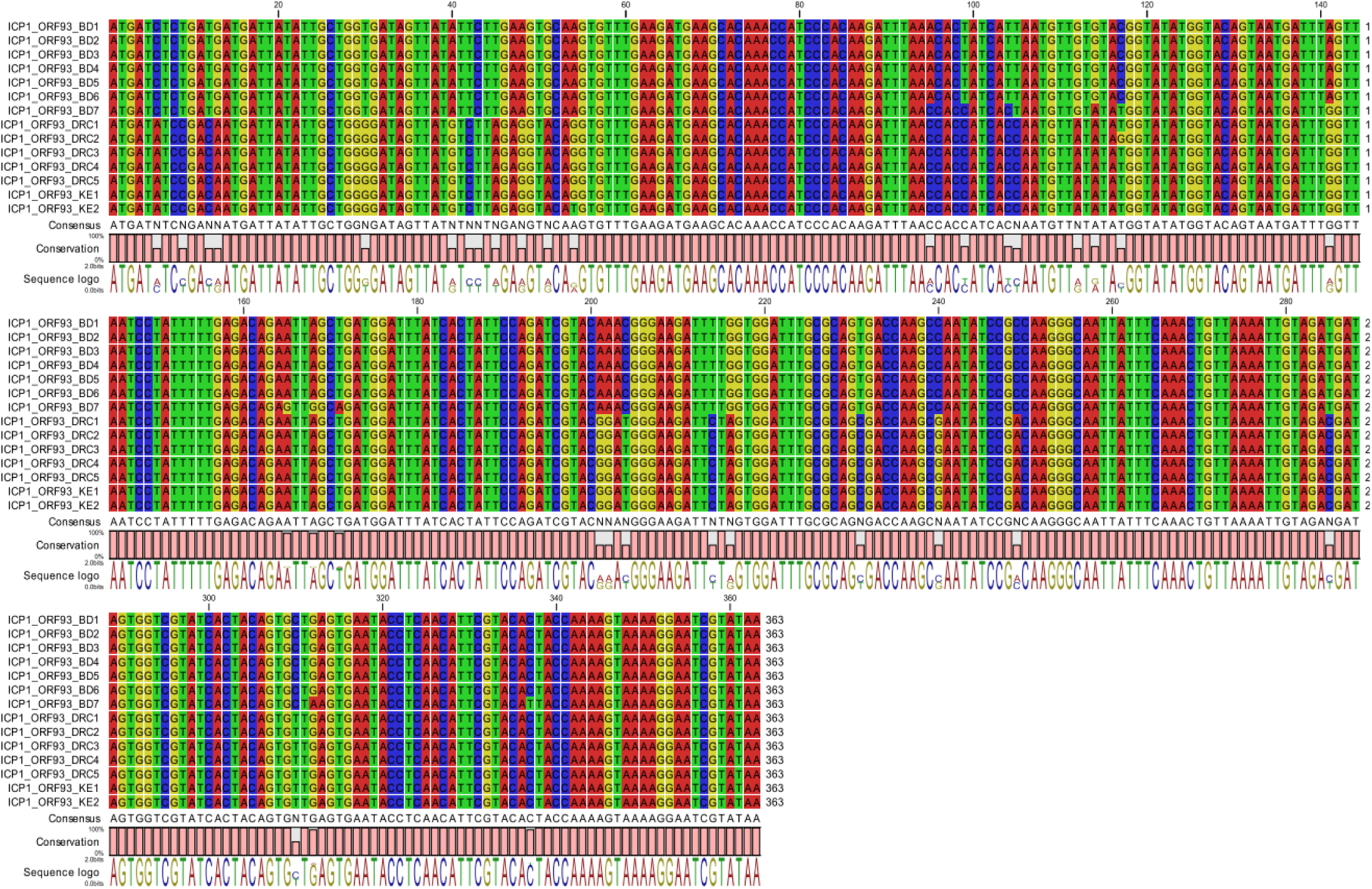
Nucleic acid alignment of ICP1 tail fiber ORF93 from Bangladesh (BD), Democratic Republic of Congo (DRC), and Kenya (KE). Sequences were obtained by PCR amplification and sequencing of clinical samples. Data visualizations prepared in QIAGEN CLC.

**Fig S4.**
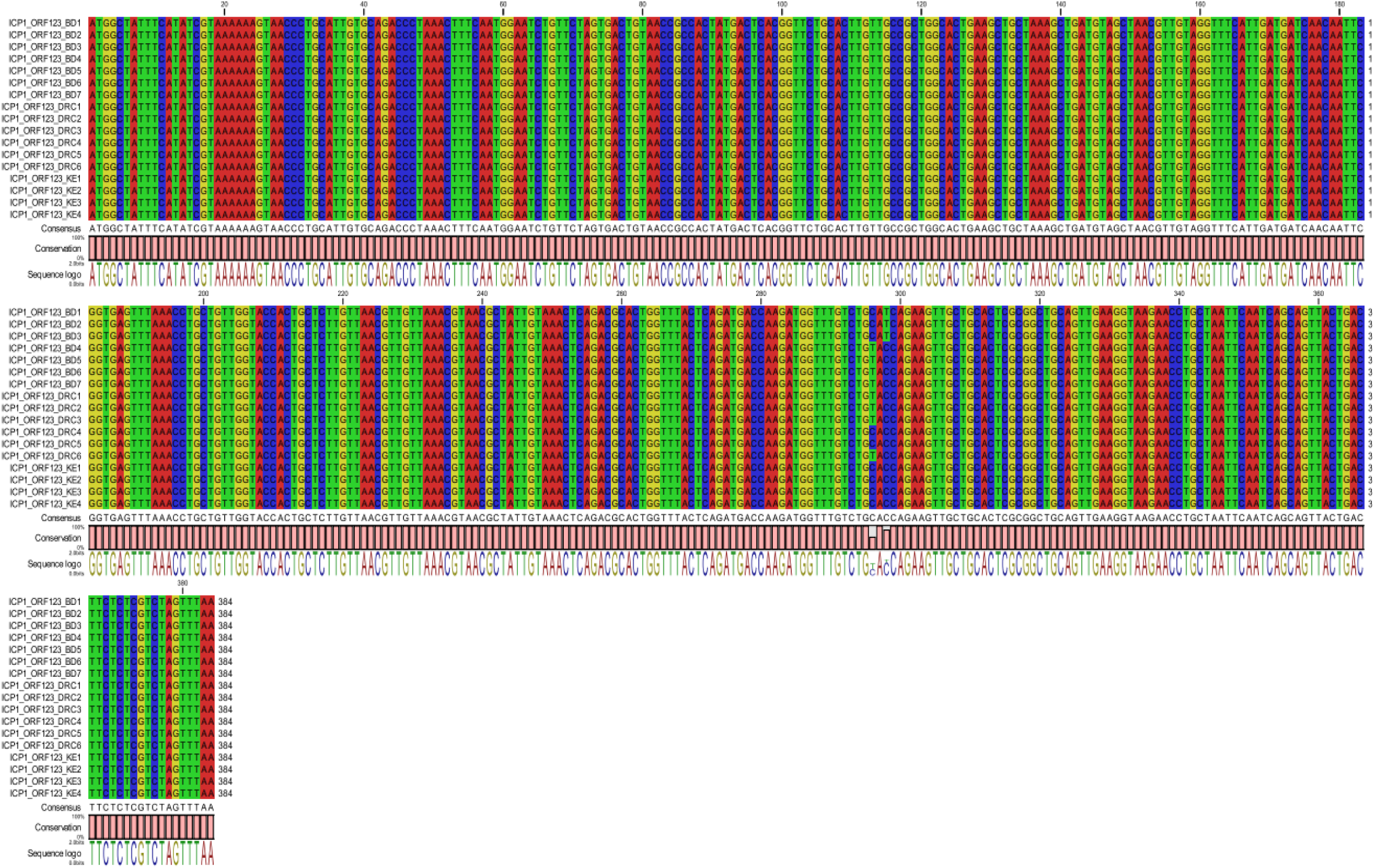
Nucleic acid alignment of ICP1 head decoration protein from Bangladesh (BD), Democratic Republic of Congo (DRC), and Kenya (KE). Sequences were obtained by PCR amplification and sequencing of clinical samples. Data visualizations prepared in QIAGEN CLC.

**Fig S5.**
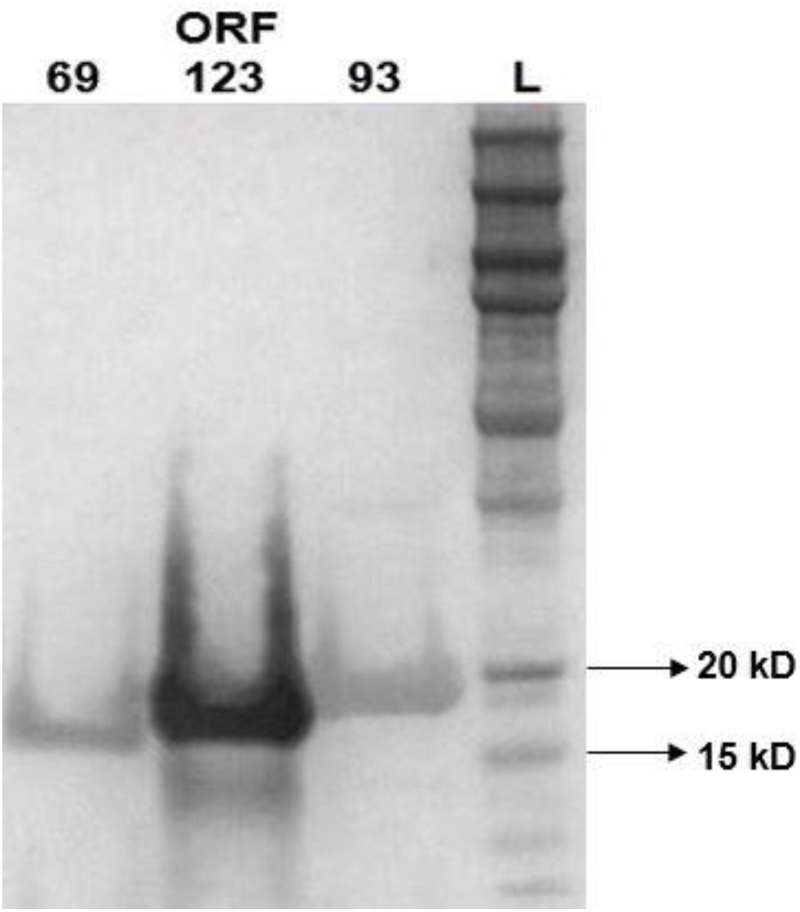
Western blot of His-tagged tail fiber ORF69, head decoration protein ORF123 and tail fiber ORF93 after purification. L = protein marker ladder.

**Fig S6.**
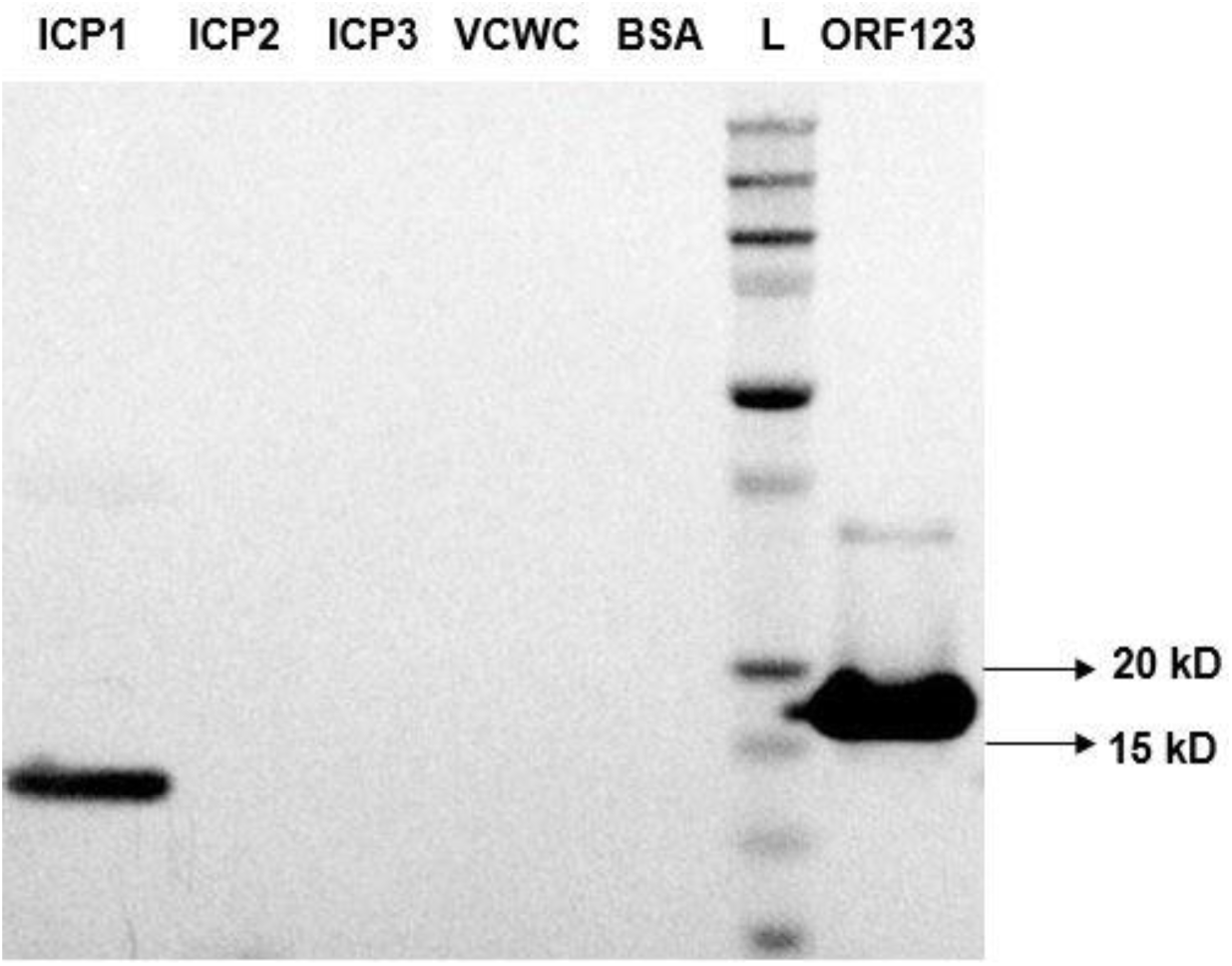
Western blot analysis of candidate ICP1 head decoration protein ORF123_mAbCL14 against ICP1. Similar results were observed for ICP1ORF123_mAbCL15 and CL16 (not shown here). Negative controls are ICP2 and ICP3. VCWC = formalin-killed *V. cholerae* whole-cell, bovine serum albumin = BSA. L = ladder (protein marker), ORF123 = ICP1 recombinant proteins (positive control).

**Fig S7.**
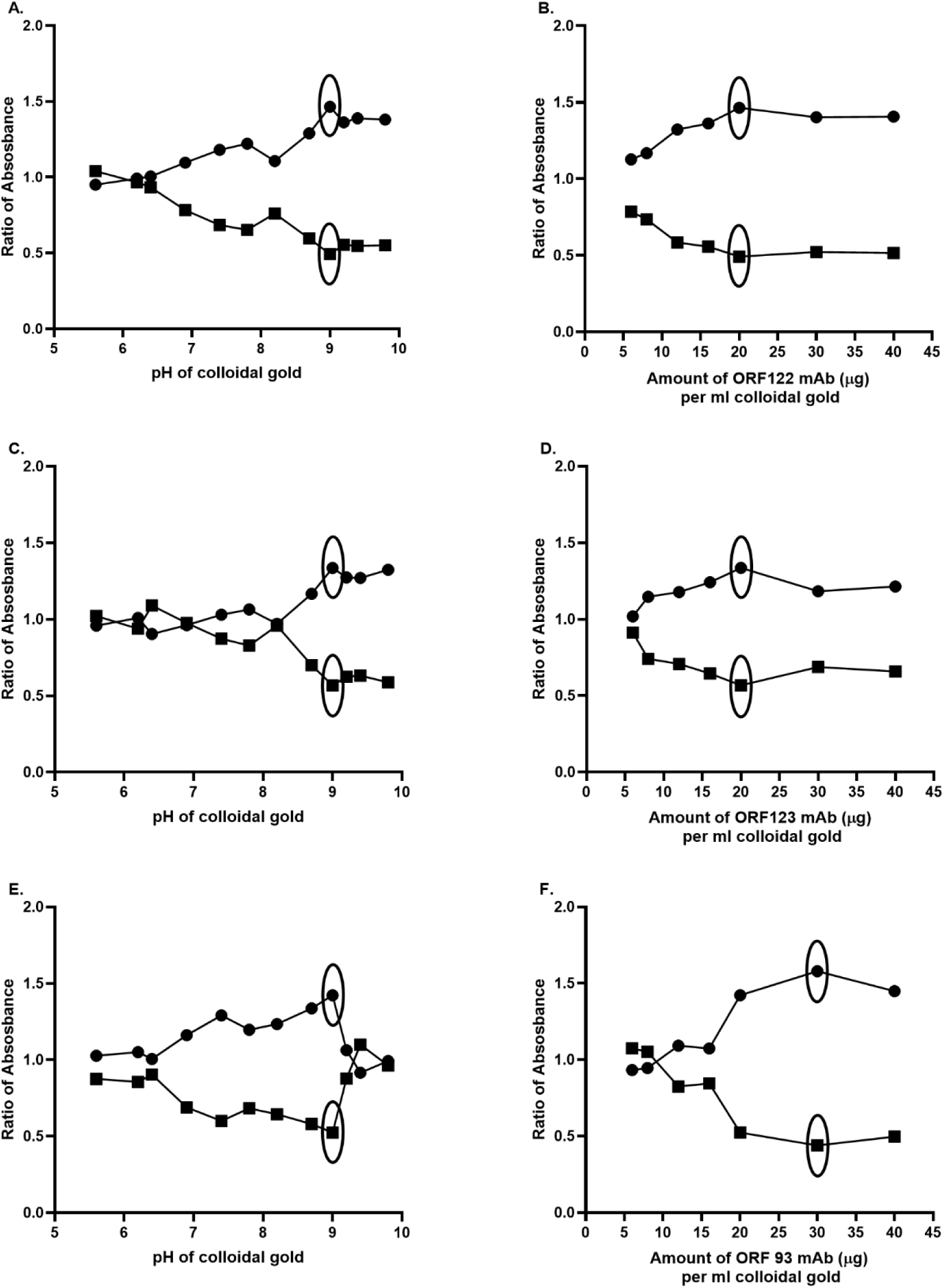
Optimization of pH and minimum mAb concentration for ORF122 mAb (A,B), ORF123 mAb (C,D) and ORF93 mAb (E,F) gold conjugation by aggregation test. Here, a ratio of absorbance at 520 nm and 580 nm represents stability, and the ratio of 600 nm to 520 indicates polydispersity of conjugated gold solution.

**Fig S8.**
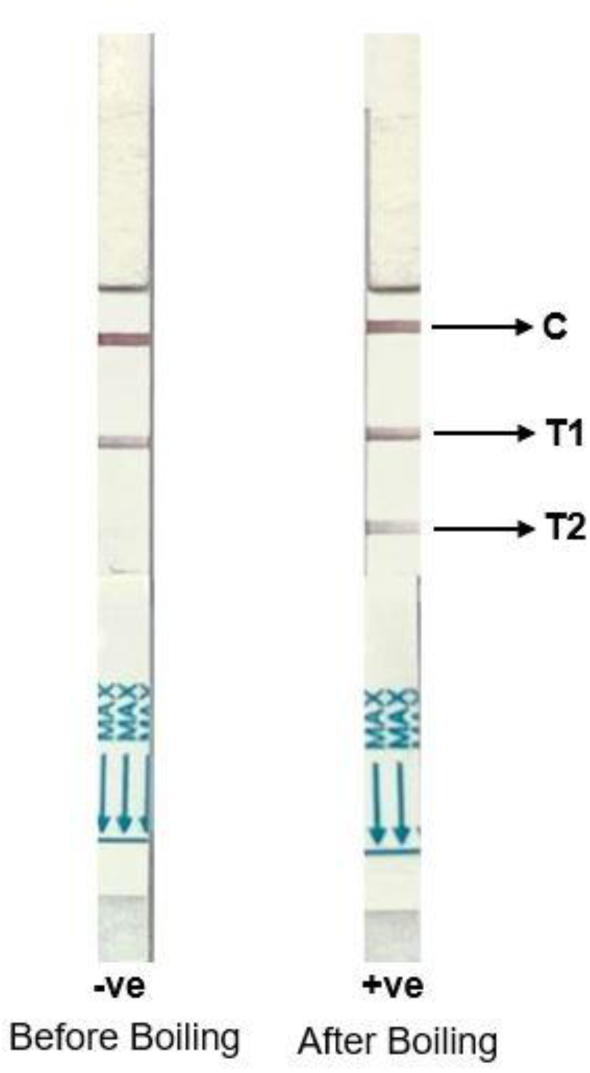
Test results of ICP1 RDTplus for the detection of both cholera LPS and ICP1. RDTplus was tested with 1:10 diluted native and boiled ICP1. Here, C = control line, T1= test line for VC LPS, T2=test line for ICP1, -ve= negative result and +ve=positive result. Red lines indicate positive control or test line. Positive C line ensures the validity of the RDT prototype result.

**Table S1.**
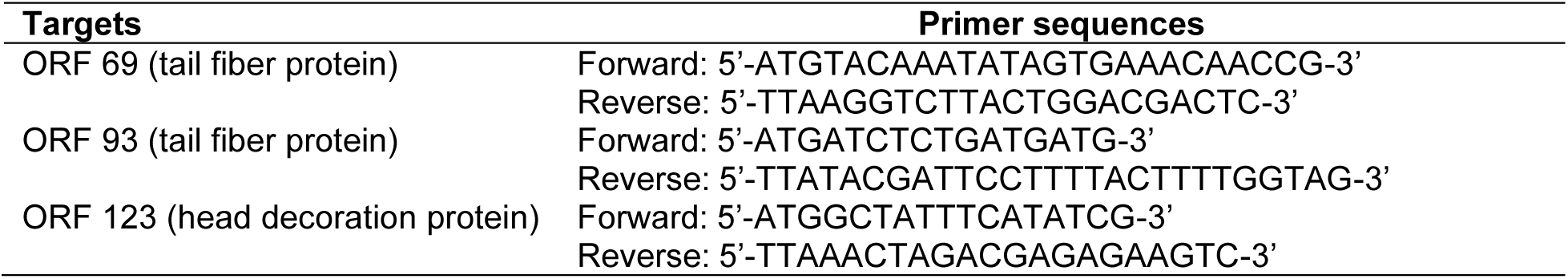
List of primers used in molecular analysis.

**Table S2.** Comparison of new target amino acid and nucleic acid sequences from Bangladesh and DRC ICP1 isolate genomic sequences (28, 29).

| Targets | Length (bp) | Mass (Da) | Nucleic Acid similarity (%) | Amino Acid similarity (%) |
| --- | --- | --- | --- | --- |
| ORF 69 (tail fiber protein) | 414 | 15,966 | 99.5 | 98.5 |
| ORF 70 (tail fiber protein) | 1068 | 38,096 | 98.9 | 97.0 |
| ORF 84 (tail sheath protein) | 1392 | 50,897 | 91.7 | 95.0 |
| ORF 93 (tail fiber protein) | 363 | 13,537 | 92.6 | 96.0 |
| ORF 123 (head decoration protein) | 384 | 13,193 | 99.7 | 99.2 |

**Table S3.** Comparison of new target amino acid and nucleic acid sequences obtained by PCR and sequencing of ICP1 positive stool samples in Bangladesh, DRC and Kenya.

| Targets | Length (bp) | Mass (Da) | Nucleic Acid similarity (%) | Amino Acid similarity (%) |
| --- | --- | --- | --- | --- |
| ORF 69 (tail fiber protein) | 414 | 15,966 | 99.5-100 | 98.5-100 |
| ORF 93 (tail fiber protein) | 363 | 13,537 | 90.4-100 | 94-100 |
| ORF 123 (head decoration protein) | 384 | 13,193 | 99.4-100 | 98.4-100 |

